# Phylogeny-aware comparative genomics of *Vibrio vulnificus* links genetic traits to pathogenicity

**DOI:** 10.1101/2025.06.02.657354

**Authors:** Luis F. Delgado, David J. Riedinger, Victor Fernández-Juárez, Daniel P. R. Herlemann, Christian Pansch, Marija Kataržytė, Greta Gyraitė, Thorsten B. H. Reusch, Marcin Rakowski, Kasia Piwosz, Adam Woźniczka, Heike Benterbusch-Brockmöller, Theodor Sperlea, Susann Dupke, Holger C. Scholz, Sandra Kube, Lasse Riemann, Matthias Labrenz, Anders F. Andersson

## Abstract

*Vibrio vulnificus* is a natural inhabitant of coastal brackish waters worldwide and an opportunistic pathogen that can cause severe infections and septicemia through seafood consumption or wound exposure. Due to global warming, its abundance is increasing at high latitudes. While the species harbors diverse virulence factors, its precise disease mechanisms remain unclear. Comparative genomics between clinical and environmental isolates can help identify key virulence genes, but the limited availability of genomes from environmental isolates has hindered progress. In this study, we sequenced the genomes of 82 *V. vulnificus* isolates from water, sediment, and seagrass along the Baltic Sea coast and complemented with published genomes from 208 clinical and 117 globally distributed isolates for comparative analysis. Phylogenetic reconstruction confirmed four major lineages, with Baltic Sea strains confined to lineage L2 and L4, while clinical and environmental strains were distributed across all lineages. This suggests that the phylogenetic structure of *V. vulnificus* reflects adaptation to environmental conditions rather than pathogenicity. Using the PhyloBOTL pipeline developed here, we identified 128 orthologs significantly enriched in clinical isolates, grouped into 36 co-localization clusters based on proximity in the genomes. These included genes linked to virulence, such as those for capsular polysaccharide synthesis and biofilm formation, as well as previously unrecognized candidates, including chaperone-usher pilus biosynthesis, spermidine synthesis, Type VI secretion effectors, and an RTX toxin-like protein. Several of the clinically enriched gene clusters have been independently lost in three *V. vulnificus* clades, suggesting convergent evolution and a distinct ecological niche shared by these claded. Finally, we used the clinically enriched genes to design PCR primers for detecting and monitoring pathogenic *V. vulnificus* strains, providing a valuable tool for surveillance and public health efforts.

## Introduction

*Vibrio vulnificus*, a Gram-negative, rod-shaped bacterium within the *Vibrio* genus, naturally inhabits estuarine, coastal, and brackish waters [1]. This opportunistic pathogen, capable of causing severe and occasionally lethal infections, poses a significant threat to humans and is a prominent contributor to non-cholera *Vibrio* related fatalities globally [2].

*V. vulnificus* can gain entry into the human body through open wounds in the skin, with even minor wounds serving as potential points of entry [3]. This can result in severe wound infections that may necessitate limb amputation and can progress to an overwhelming septic shock, frequently culminating in fatality due to multiple organ failure [4]. Another transmission route is through the consumption of raw contaminated seafood, particularly oysters [5]. The most susceptible populations for fulminant extraintestinal infections from either the oral or cutaneous route are individuals with compromised immune responses. This includes patients with thalassemia, diabetes, HIV, or liver diseases (cirrhosis or hepatitis) and those receiving immunosuppressant drugs for other underlying conditions [6,7]. A comprehensive review of 459 U.S. cases reported to the Food and Drug Administration (FDA) between 1992 and 2007 revealed a staggering 51.6% mortality rate among patients infected with *V. vulnificus* [8].

Infections associated with *Vibrio* spp. exhibit a correlation with elevated water temperatures surpassing approximately 15°C and a moderate salinity range of PSU 2 -25 [9]. Over the past decade, in tandem with the warming trends observed in the North and Baltic Seas, there has been a surge in reported cases of *Vibrio* infections in this region [10–12]. Recent surveys of *V. vulnificus* in coastal zones of the Baltic Sea have confirmed the regulatory influence of temperature and salinity on the species, while also suggesting a potential role for eutrophication status [13,14]. Since the trend of increasing *Vibrio*-related infections and fatalities along the Baltic Sea coasts is expected to be worsened by climate change [15], thereby posing a significant threat to human health and tourism industry, assays for fast and accurate detection of *V. vulnificus* in environmental as well as seafood samples, are needed [16].

*V. vulnificus* harbors a wide array of virulence factors, including mechanisms for acid neutralization, capsular polysaccharide production, iron acquisition, cytotoxicity, motility and adhesion [8]. However, its precise disease mechanisms remain poorly defined. Mouse model studies have shown that not all *V. vulnificus* isolates have the same virulence potential [17,18]. However, unlike *Vibrio cholerae*, where all cholera-causing strains are affiliated with a single clade [19], genomic analyses of *V. vulnificus* have revealed that clinical strains exhibit extensive genomic diversity. Comparative genomics of clinical and environmental isolates makes it possible to identify key virulence factors and genomic signatures associated with pathogenicity. Since the first *V. vulnificus* genome (CMCP6) was sequenced in 2002 [20], more than 500 genomes have been made publicly available in GenBank—approximately 40% from clinical isolates and 60% from environmental sources. However, the majority of environmental isolates originate from animal reservoirs—primarily fish and shellfish. Only a limited number are derived directly from environmental matrices such as water and sediments, where selection pressure for maintaining virulence genes is likely lower than in host-associated environments. While the growing availability of genome sequences has advanced our understanding of *V. vulnificus* population structure [21], phylogenetics [22,23], and virulence [24–26], key aspects of its pathogenic mechanisms remain unresolved. Notably, the limited representation of genomes from non-animal, environmental sources has remained a gap in current genomic resources.

Here, we performed whole-genome sequencing and assembly of 82 environmental strains of *V. vulnificus* isolated from water, sediment and seagrass along the Baltic Sea coast. These genomes provide insights into the species’ genomic diversity in this region. By integrating them with publicly available genomes from environmental and clinical sources worldwide, and applying the PhyloBOTL (Phylogeny-Based Ortholog-Trait Linkage) pipeline developed in this study, we link gene functions to pathogenicity and identify a number of candidate virulence genes. Finally, we design novel PCR primers targeting genes enriched in clinical isolates to support environmental surveillance of *V. vulnificus*.

## Results and discussion

### Phylogenetic structure of *V. vulnificus*

We sequenced the genomes of 82 *V. vulnificus* strains isolated from five different sites along the Baltic Sea coastline during the summer of 2021 [Citation error]. These sites encompassed two locations in Sweden, one in Poland and two in Germany (Fig 1 and S1 File). The water temperature and salinity at the sites ranged from 16.5 −20.3℃ and 6.5 −10.4 PSU, respectively (Table 1). The strains were isolated from the water column, from the top sediment layer, and from eelgrass (*Zostera marina*) leaves, within or proximate to eelgrass meadows (S1 Table provides contextual data for the isolates). DNA sequencing and assembly yielded genomes ranging in size from 4,838,252 to 5,241,188 base pairs (bp) (S2 Table). Seventeen of the strains harbored plasmids, ranging in size from 44,678 to 143,195 bp, and 53 were predicted to have prophage regions, ranging from 2101 to 120,375 bp. Several antibiotic resistance genes were detected in the 82 isolates. Among these, the genes varG and tet(34) are present in all, and tet(35) is only absent in six isolates. These genes confer resistance to various antibiotic classes including carbapenem, tetracycline and beta-lactam (S3 Table).

**Fig 1.**
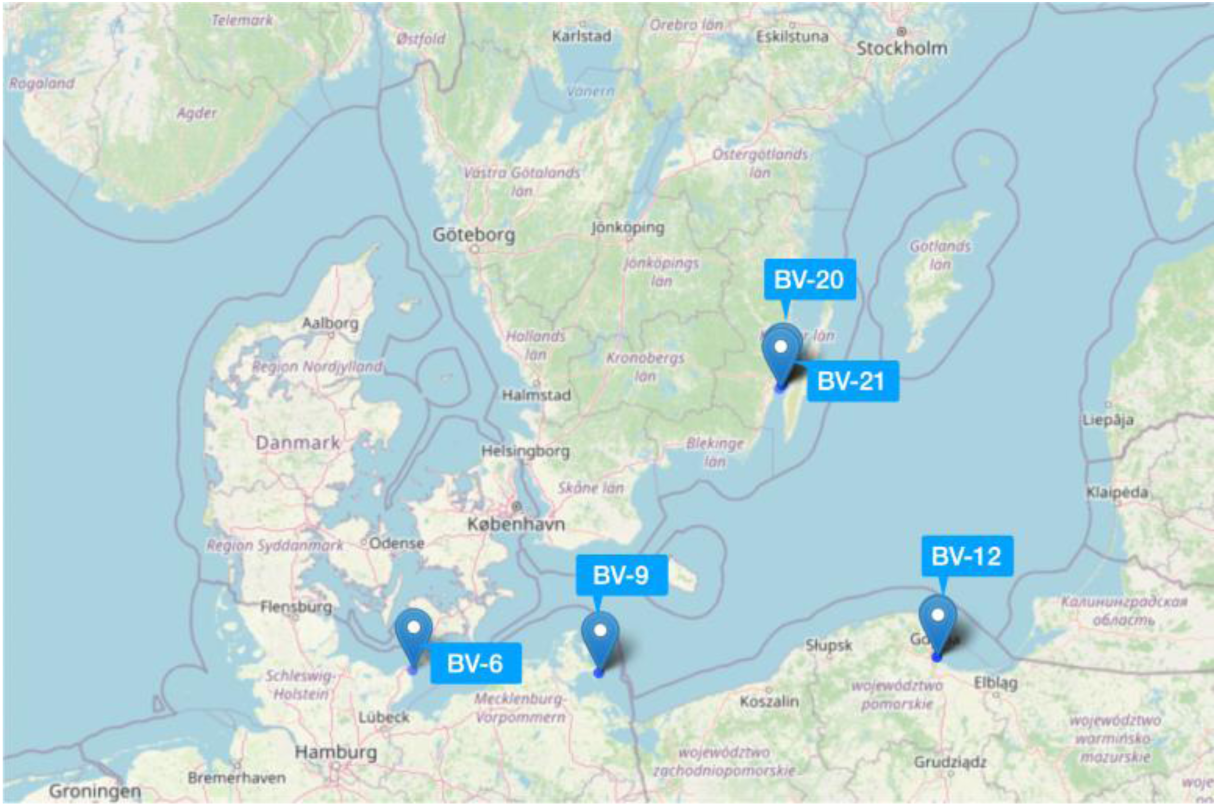
Sampling locations of the *Vibrio vulnificus* isolates sequenced in this study. ‘BV’ denotes BaltVib, the larger project this study was part of.

**Table 1.** Sampling stations of the *V. vulnificus* isolates sequenced in this study. Isolation source: w = water; s = sediment; z = *Zostera marina* leaf. Additional contextual data is provided in S1 Table.

| Station | No. isolates | Country | Average water temperature (°C) | Average water salinity (PSU) |
| --- | --- | --- | --- | --- |
| BV-21 | 1(w) | Sweden | 16.5 | 6.7 |
| BV-20 | 11(w); 9(s) | Sweden | 16.8 | 6.5 |
| BV-12 | 6(w); 47(s); 2(z) | Poland | 20 | 7.0 |
| BV-09 | 1(w); 2(s) | Germany | 20.3 | 7.7 |
| BV-06 | 3(s) | Germany | 19.3 | 10.5 |

We complemented the 82 new genomes with 325 previously published environmental (n=117) and clinical (n=208) *V. vulnificus* genomes to facilitate a comprehensive comparative analysis (S2 Table provides metadata on the included genomes). Of the previously published genomes, four of the environmental and 47 of the clinical (mainly from the 2018 heatwave [10,11]) stem from the Baltic Sea region (Germany, Denmark, Norway, Sweden, Finland). The remaining genomes are primarily from the USA (n=204), China (n=26), Israel (n=10), South Korea (n=9) and Colombia (n=9). We used maximum-likelihood to infer the phylogeny of the 407 strains, leveraging an alignment of 2,223,636 nucleotide positions from the core genome, with 361,414 positions exhibiting variation. Consistent with the findings of Roig et al. [22] and subsequently Lopez et al. [25], our study confirmed the formation of four major lineages (L1 -L4) among the strains. Additionally, consistent with Roig et al., we identified a fifth lineage (L5) comprising a single genome. Furthermore, our expanded dataset revealed three additional lineages, encompassing six genomes in total, positioned between L5 and L2 (Fig 2).

**Fig 2.**
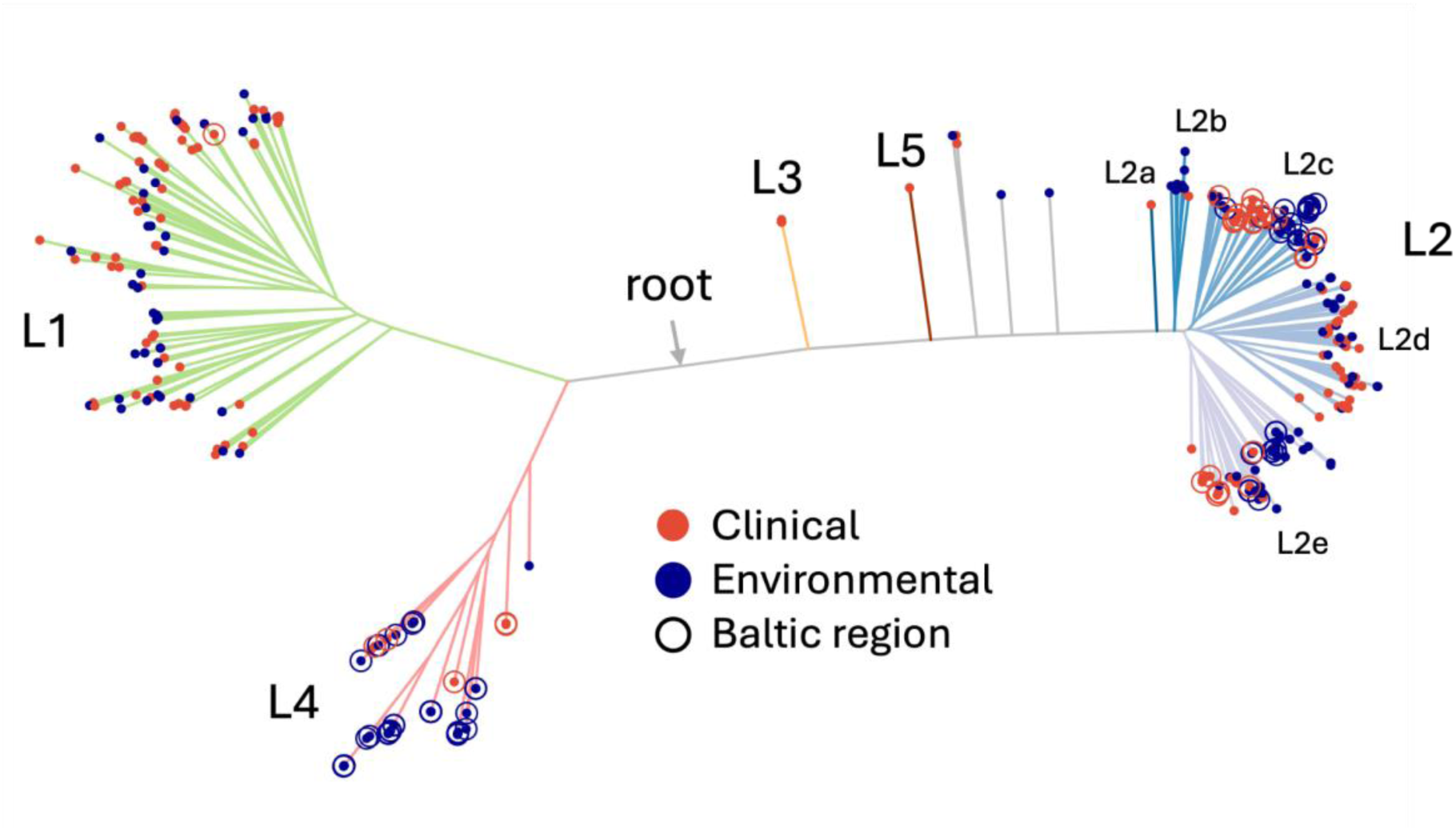
Maximum-likelihood phylogeny based on protein-coding regions of the core genome. Lineage numbers are according to Roig et al. [22], but the L2 sublineages (L2a - L2e) were introduced in this study. Branches are colored according to (sub)lineage. Leaf colors indicate clinical (n=208) or environmental (n=199) isolate source and circles indicate Baltic Sea region isolates (n=133, of which 82 were sequenced in this study). The approximate location of the root is indicated by the arrow and was determined in a separate phylogenetic reconstruction that also included other *Vibrio* species.

Notably, all of the new 82 Baltic Sea genomes fall in the L2 and L4 lineages, which is also the case for the previously published 51 genomes from the Baltic Sea region (except for a single isolate; DSM 11507; for which the isolation source is unclear). Lineage L4 comprises a small number of previously published isolates from Germany and Sweden and two isolates from the Ebro Delta in Spain [22]. The lineage is now greatly expanded, comprising 46 genomes from Sweden, Poland and Germany. Lineage L2 is composed of five sublineages, which we denote L2a -L2e. Interestingly, the Baltic Sea region genomes are confined to sublineages L2c and L2e, collectively representing 61% of the genomes in these sublineages (Fig 2).

The biased phylogenetic distribution of the Baltic Sea region genomes may suggest that the phylogenetic lineages of *V. vulnificus* constitute strains adapted to distinct environmental conditions. Temperature emerges as a plausible factor, since the cold winters in the Baltic Sea region may require specific adaptations not needed at lower latitudes with warmer climate. However, the presence of multiple isolates from tropical regions in L2c and L2e contradicts the notion of these sublineages being exclusively cold-adapted. Salinity is another potential factor. In the central to southwestern Baltic Sea, salinity ranges from 6 -14 PSU, while *V. vulnificus* can thrive in environments with considerably higher salinity. Due to the unavailability of data on salinity, temperature, and other physicochemical parameters for most published environmental genomes, a comprehensive phylogenomic assessment of these factors remains unfeasible at present. However, a recent study by Lopez et al. [25] of two sites in Florida indicates the importance of salinity. Specifically, six out of seven isolates from the site with higher salinity (29 - 32 PSU) clustered within lineage L1, whereas 11 out of 14 isolates from the site with lower salinity (6 - 19 PSU) clustered within L2. There was no significant temperature difference between the sites, so salinity appears more important in this case. However, the underlying drivers may be other environmental factors differing between the sites.

Since most human clinical isolates have earlier been found to cluster in lineage L1, this lineage has been proposed to carry an increased genetic potential for virulence [25]. However, this view was recently challenged by a study sequencing a large number of clinical isolates from the US, where the isolates were found to be equally distributed between L1 and L2, although infections caused by L1 isolates demonstrated a borderline significant higher mortality [25]. Our finding that both clinical and environmental isolates from the Baltic Sea region are confined to Lineage L2 and L4 reinforces that clinical strains are not restricted to L1 but instead suggests the phylogenetic structure of the species reflects environmental factors. Another potential explanation could be geographic barrier effects - i.e. that the sublineages are restricted to geographic regions due to dispersal limitation [21].

### Genes enriched in clinical isolates

The expanded number of genomes from environmental strains allows identifying genes associated with pathogenicity by comparing the gene content in the environmental and clinical isolates. Although many of the environmental isolates may well have pathogenic potential we can assume that a fraction of them lack such capacity, and thus genes that are significantly enriched in the clinical isolates are potentially of importance for the pathogenicity trait. We used a phylogeny-aware approach [27] that identifies genes (orthologs) whose presence co-evolves with a trait over the phylogenetic tree. Thus, orthologs that are gained (or lost) in conjunction with switching from the environmental to clinical “trait” more often than expected by chance over the evolution will be identified. We identified 13,920 orthologous groups of genes (OGs, also sometimes referred to as ‘orthologs’ in the following) among the 407 genomes. Among these, 128 were found to be enriched and 82 depleted in the clinical vs. environmental strains (false discovery rate-adjusted *P*-value < 0.05). The 128 clinically enriched orthologs had significantly more matches in the Virulence Factor Database [28] than orthologs found in at least 95% of the genomes (Chi-squared test, *P*-value = 0.0035) supporting the strategy. In the following sections we will focus on the orthologs enriched in the clinical strains.

Since functionally related genes are often encoded in the same genomic region on bacterial chromosomes [29,30], we used information on the orthologs’ pairwise proximity in the *V. vulnificus* genomes to cluster them (see Materials and methods). This resulted in 36 co-localisation clusters of which 20 were singletons and the remaining consisted of 2 to 24 OGs (Fig 3 and 4; Table 2; detailed information in S4 Table). Twenty-one of these were located on chromosome I, fourteen on chromosome II, and one on a plasmid. The co-localisation clusters entailed both those containing genes previously linked to pathogenicity in *V. vulnificus,* such as genes for biofilm formation and capsular polysaccharide synthesis, and clusters with genes that - to our knowledge - have not previously been associated with pathogenicity in the species. These include genes of the chaperone-usher pathway for pilus synthesis, genes for spermidine synthesis, genes encoding effector proteins of the Type VI secretion system, and a gene for an RTX toxin-like protein. In the following sections, we provide a more detailed description of selected co-localization clusters, with the aim of encouraging follow-up studies to further investigate their potential roles in pathogenicity. In addition to our internal OG IDs, gene IDs for the corresponding genes in the CMC6 strain are provided when the gene is present.

**Fig 3.**
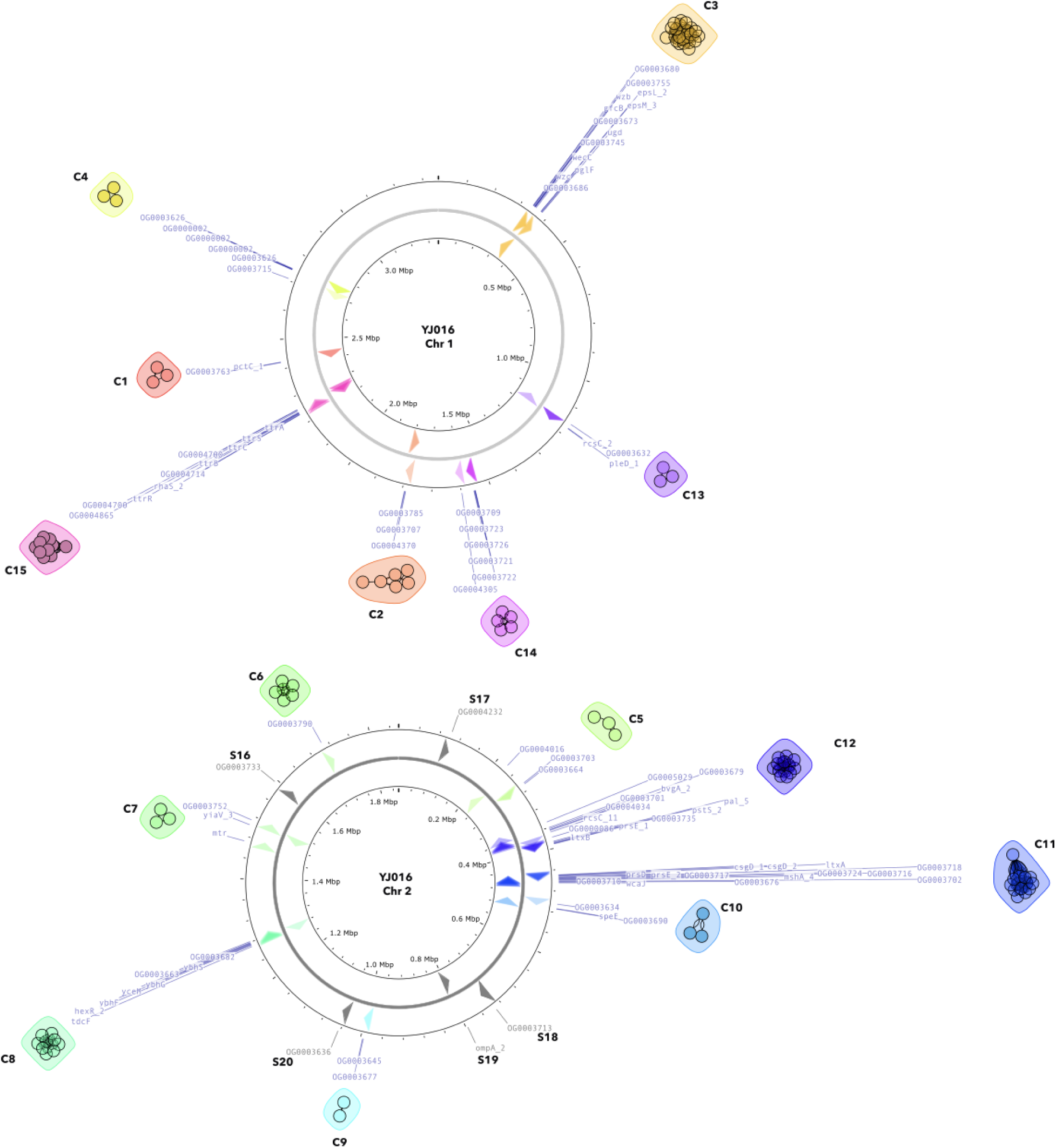
Co-localisation clusters of clinically enriched orthologs. The orthologs’ positions and directionality in the two *V. vulnificus* YJ016 chromosomes are indicated with arrows, colored according to co-localisation clusters. Gray arrows represent singleton clusters. Not all singleton clusters are present in strain YJ016.

**Fig 4.**
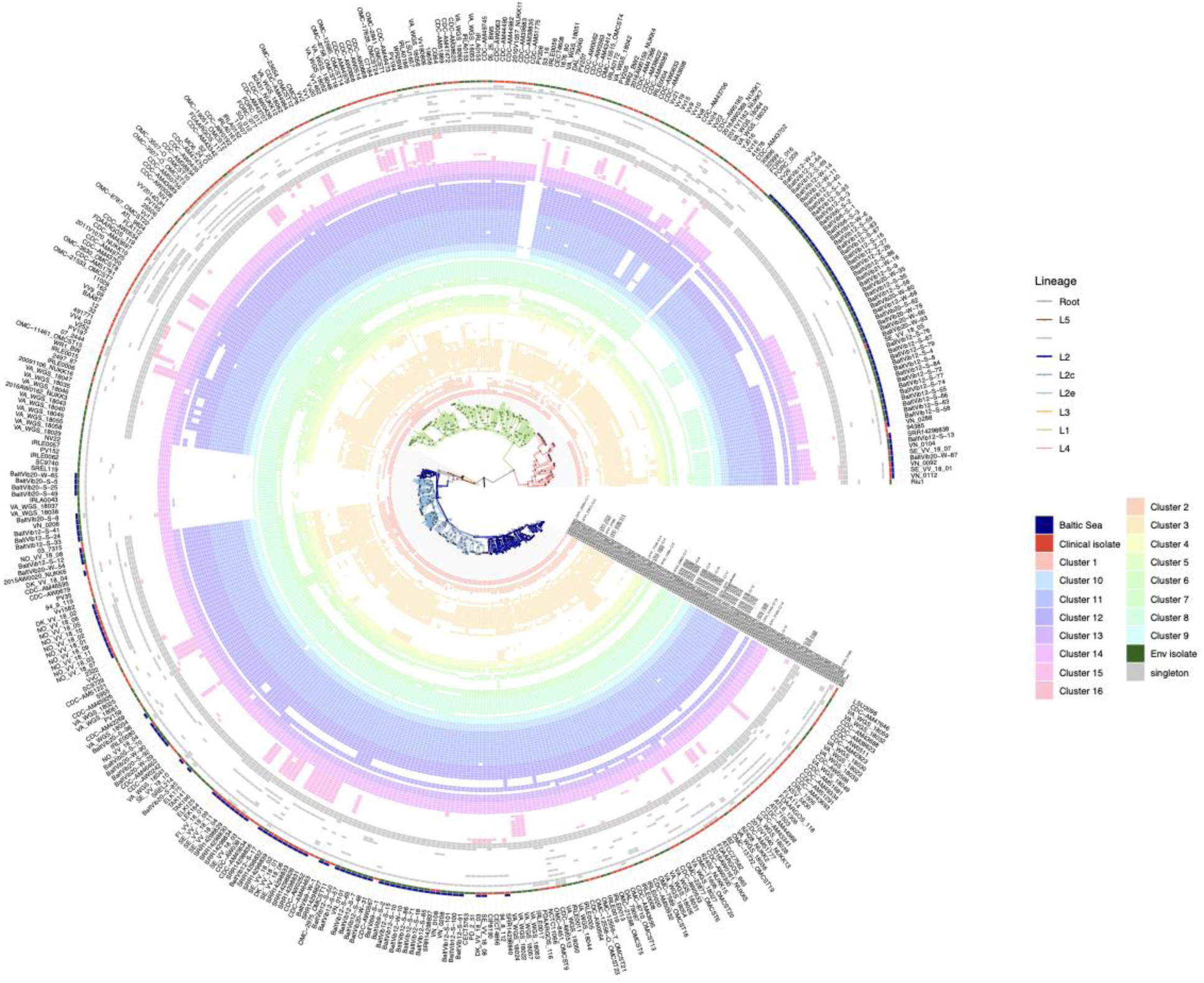
Presence of the 128 clinically enriched orthologs in the 407 *V. vulnificus* genomes. The orthologs (one per row) are ordered and colored according to co-localisation clusters. The outer colored rings indicate if the isolate is from the Baltic Sea region and if it is a clinical isolate. The branches of the phylogenetic tree are colored according to (sub)lineage.

**Table 2.**
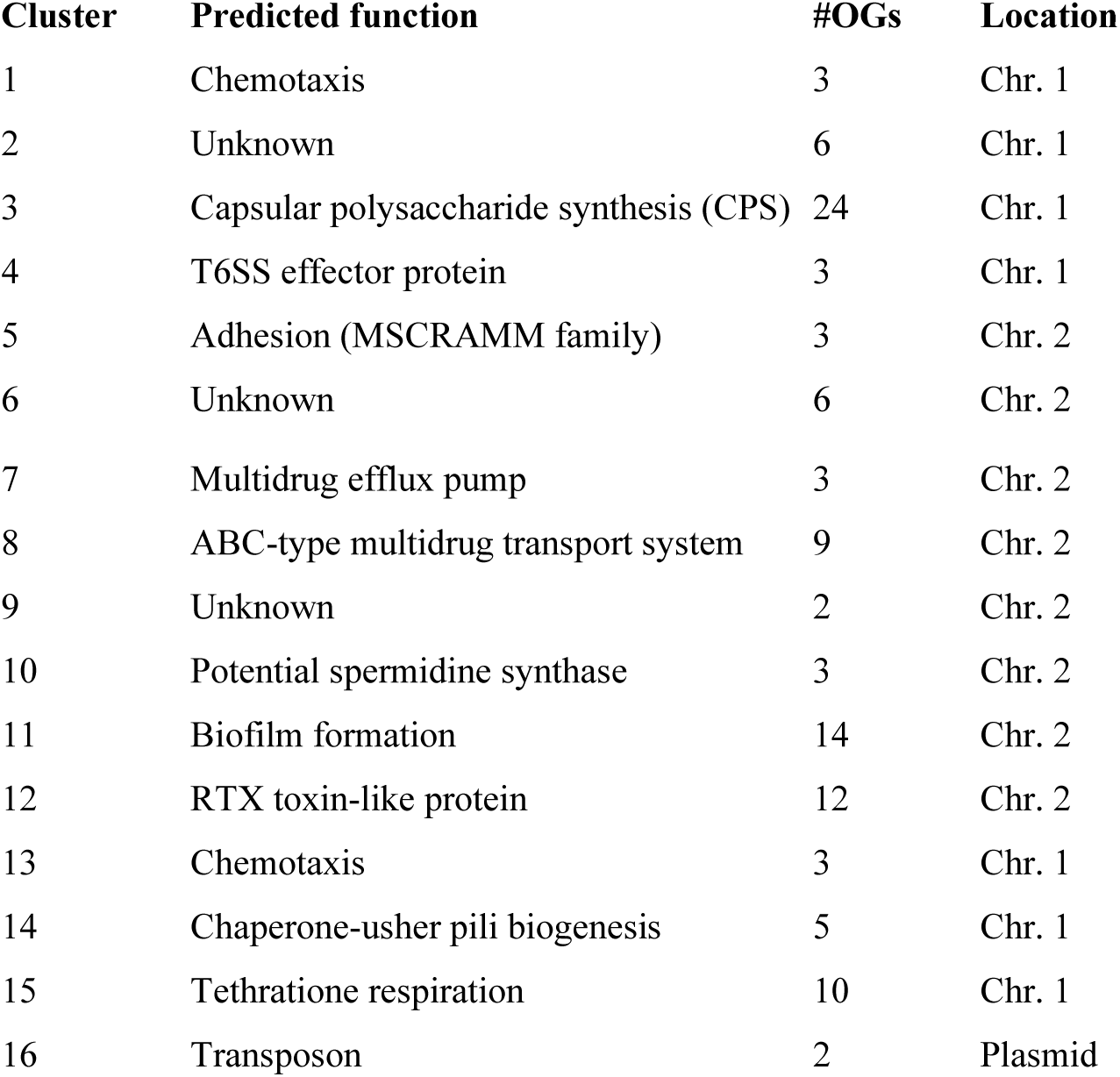
Summary of co-localisation clusters of clinically enriched orthologs. Predicted function refers to the function of the cluster as a whole or of an ortholog within the cluster. Only clusters with >1 ortholog are shown. See S4 Tab for information on all clusters and individual orthologs.

Cluster 3 is the largest cluster, containing 24 enriched orthologs, located on chromosome I. This cluster includes several genes known to be involved in capsular polysaccharide (CPS) synthesis [31]. CPS is considered the most important virulence factor of *V. vulnificus* [32]. It mediates resistance to complement-mediated bacteriolysis and phagocytosis, and is essential for the bacterium’s survival in human serum [33]. Its expression has been demonstrated to undergo phase variation [34,35], and acapsular strains have proven to be non-infectious [36]. CPS is not required for biofilm formation in *V. vulnificus* [37] and its expression even inhibits biofilm formation [38]. The CPS synthesis gene locus of *V. vulnificus* is similar to that of the group 1 or 4 capsules of *Escherichia coli* [39]. As shown in S1 Fig, the cluster is highly diverse, with the enriched orthologs found in 322 different gene order configurations, and never more than six isolates sharing the same configuration. Eleven of the orthologs are more prevalent than the other in the cluster, present in 92-100% of the clinical and 70-88% of the environmental strains. Among these are *wza-wzb-wzc*, responsible for polymerization control and translocation of CPS across the outer membrane [31], all of which are present in 352 out of 407 strains. Of the 55 strains where one or more is missing, only five are clinical, in line with CPS being a crucial virulence factor.

Cluster 11 comprises 14 orthologs situated in a genomic region on chromosome II involved in formation of biofilm. The enriched orthologs include the genes of the *cabABC* operon, responsible for production and secretion of the calcium-binding extracellular matrix protein CabA, the *brpABCDFHIJK* locus, which is one of several loci responsible for assembly and export of exopolysaccharide (EPS) in *V. vulnificus*, and the regulatory genes *brpT* and *brpS* [40,41] (S2 Fig). While EPS is the predominant component of the *V. vulnificus* biofilm matrix, CabA contributes to the structural integrity and robustness of the biofilms [42]. The expression of *cabABC* and the *brp* locus undergoes regulation through a complex regulatory network involving BrpS, BrpT, and the master regulator BrpR, and is activated under conditions of elevated levels of the intracellular second messenger cyclic dimeric guanosine monophosphate (c-di-GMP) [37,40,43,44]. Although *V. vulnificus* encodes nearly 100 proteins predicted to synthesize, degrade, and bind c-di-GMP, relatively little is known about what environmental signals regulate c-di-GMP levels, and thereby biofilm formation [45]. However, calcium was shown to be a primary environmental signal that specifically increases intracellular c-di-GMP concentrations, which in turn triggers biofilm formation [45]. The calcium-induced biofilm formation is likely important for colonization and accumulation of *V. vulnificus* in oysters and other bivalves that have high concentrations of calcium on their surfaces [46].

Cluster S16 includes a single ortholog (OG0003733; VV2_1694) that is functionally related to Cluster 11. The ortholog corresponds to *brpN,* a acyltransferase-3 domain-harboring protein recently found to be important for EPS production in *V. vulnificus* [47]. Together with the above described *cabABC* and *brpABCDFHIJK* loci, as well as a third locus for EPS synthesis - the *brpLG* locus [37,40,43,44] - *brpN* is essential for the formation of robust biofilms and rugose colonies in *V. vulnificus*. Similar to the other loci, it is regulated by BrpT, and ultimately by the master regulator BrpR [47].

Cluster 14 consists of five enriched orthologs, arranged as an operon on chromosome I. This operon includes a pilus assembly chaperone (OG0003726; VV1_2742), an outer membrane pilus assembly protein (usher) (OG0003709; VV1_2741), a fimbrial/pilus subunit protein (OG0003723; VV1_2740), a LuxR-family response regulator (OG0003721; VV1_2744), and a protein with an unknown function (OG0003722; VV1_2743) (S3 Fig). Alongside another operon in *V. vulnificus*, it has been categorized within the archaic σ-clade of the chaperone–usher (CU) pathway for pilus biogenesis [48]. Although widely distributed among Gram-negative bacteria, the structural makeup of these archaic pili has remained elusive, until a recent study revealed their assembly into an ultrathin, superelastic spring structure [49]. In *Acinetobacter baumannii* and *Pseudomonas aeruginosa*, σ-clade pili play a role in surface attachment and biofilm formation [50,51]. While the involvement of multiple Type IV pilus systems in *V. vulnificus* is well-documented, contributing to biofilm formation, adherence to epithelial cells, and virulence [52,53], the ecological significance and potential virulence of σ-clade pili, to our knowledge, remain unexplored in the species. However, in a gene expression study comparing normal and elevated c-di-GMP conditions, the fimbrial/pilus subunit gene (VV1_2744) was repressed in elevated c-di-GMP conditions [40], indicating that these pili are not involved in biofilm formation.

Cluster 4 includes three enriched orthologs on chromosome I: a diguanylate cyclase (OG0003715; VV1_1733), a PAAR repeat-containing protein (OG0003626; VV1_1702), and a protein annotated as permease in UniRef (OG0000002; VV1_1704). Diguanylate cyclase (DGC) synthesizes c-di-GMP. In most isolates, the DGC is encoded rather distant from the other two orthologs (S4 Fig), and may not be directly functionally coupled to these. In contrast, the other two are mostly encoded adjacently, with OG0003626 upstream of OG0000002, sometimes with multiple copies of OG0000002 in series. OG0003626 encodes an N-terminal PAAR repeat followed by a MIX_IV signal. The MIX (Marker for type sIX) is commonly found in Type VI secretion system (T6SS) effector proteins that carry C-terminal toxins [54]. T6SS are commonly found in Gram-negative bacterial genomes and are made up of 13 conserved proteins. T6SS has dual functions, secreting both anti-eukaryotic and anti-prokaryotic effectors. The PAAR repeat of OG0003626 provides binding to the VgrG protein that forms the tip of the T6SS inner tube that is translocated into the prey cell [55]. The C-terminal of the protein belongs to the (CDD) BTH_I2691 family effector. This has a predicted structure similar to colicin Ia [56], a bacteriocin that kills Gram-negative bacteria by forming a pore in their inner membrane [57]. Since antibacterial pore-forming MIX effector proteins are typically encoded upstream of cognate immunity protein(s) that protect the cell from autolysis [58], it is plausible that OG0000002 encodes such immunity proteins. *V. vulnificus* is known to encode two T6SSs; T6SS2 is found in all while T6SS1 only in some isolates [59]. Both of these have been shown to exert both intra- and inter-species antibacterial activity and are believed to provide the host cell with a competitive advantage [59,60]. Similar to many other MIX effector proteins, the Cluster 4 effector protein is not encoded within the T6SS loci, but probably uses one or both of them for its secretion. Interestingly, the promoter region of OG0003626 has a binding site for SmcR [61], a quorum-sensing regulator and homolog of LuxR in *Vibrio fischeri* [62].

Cluster 12 consists of twelve enriched orthologs located on chromosome II (S5 Fig). Six of these are located within the same operon that is present in all clinical isolates. The operon encodes an RTX toxin-like protein (OG0000086; VV2_1514), a putative RTX toxin secretion ATP-binding protein (OG0003730; VV2_1515), a type I secretion membrane fusion protein (OG0003736; VV2_1516), a TolC family outer membrane protein (OG0003735; VV2_1517), an OmpA-like domain-containing protein (OG0003662; VV2_1518) and an ABC-type phosphate transport system protein (OG0003678; VV2_1519). The large (4654 aa) RTX toxin-like protein encodes a Ca+ stabilized adhesin repeat and the conserved FhaB domain, which is present in large exoproteins involved in heme utilization or adhesion [63]. Although the protein displays sequence similarity with a putative repeat-in-toxins (RTX) family exoprotein in *E. coli,* it has distinct features from typical RTX proteins [63]. Deletions in the gene in *V. vulnificus* YJ016 did not affect cell adherence, cytotoxicity, or virulence. Instead, the gene was shown to display increased expression under iron-limiting conditions [63]. Iron availability plays a pivotal role in the pathogenesis of *V. vulnificus* infection and growth [64]. To our knowledge, this is the first report linking this ortholog to pathogenicity of *V. vulnificus*.

Other noteworthy co-localisation clusters are Clusters 10, 15, 5 and 13. Cluster 10 encodes potential spermidine synthases. Spermidine has been shown to protect *Salmonella enterica* against ROS-mediated cytotoxicity [65] and spermidine transporters are upregulated in *V. vulnficus* grown in serum [66]. Cluster 15 includes the *ttrRSBCA* locus for tetrathionate respiration, enabling the utilization of tetrathionate as an electron acceptor and thus facilitating respiration in anaerobic environments, such as in sediments or inside the mammalian gut [89]. Cluster 5 includes a gene with an MSCRAMM family adhesin clumping factor ClfA region. MSCRAMM (microbial surface components recognizing adhesive matrix molecules) are proteins mainly described in Gram-positive bacteria that mediate attachment to host tissue; Clumping factor A (ClfA) is an important virulence factor of *Staphylococcus aureus* that binds to the blood plasma protein fibrinogen [92][93][67]. Cluster 13 includes a methyl-accepting chemotaxis protein comprising a Cache 3/Cache 2 fusion domain that serves as an extracellular sensor, binding to environmental signaling molecules [91]. The ligand of this protein in *V. vulnificus* is to our knowledge unknown. S1 Appendix and S6 Fig - S9 Fig provide more details on these clusters.

*V. vulnificus* is known for its high invasiveness and capacity to damage host cells, partly due to the expression of several exotoxins [46]. These include the multifunctional autoprocessing repeats-in-toxin (MARTX) toxin, in *V. vulnificus* named RtxA [68], Phospholipase A2 (PlpA) [69], Cytolysin/hemolysin (VvhA) [70], and Elastolytic protease (VvpE) [71]. Despite the fact that at least two of these, RtxA and PlpA, have proven to be essential for pathogenicity in *V. vulnificus,* none were detected by our method. The reason is that the orthologs representing these toxins are generally present across all strains, except for RtxA, which is absent in a clade of fourteen Baltic Sea environmental isolates and in one clinical isolate (S10 Fig). This highlights a weakness of the employed method; orthologs not displaying a difference in presence between the two sets of isolates will not be detected, although they may provide the bacteria with pathogenic potential.

### A common theme among several co-localisation clusters

Several of the co-localisation clusters are associated with c-di-GMP. This molecule plays a key role in controlling the transition from a motile to a sessile lifestyle. Like several other pathogenic bacteria, *V. vulnificus* can exist in a free-living planktonic state or a surface-attached biofilm state [72]. Biofilms provide resistance to environmental stressors, and in the case of *V. vulnificus*, a means for colonization and accumulation in shellfish. Biofilm formation is triggered by external stimuli, for example high calcium levels, via c-di–GMP [45]. At least two of the enriched clusters (11 and S16) encode genes related to biofilm formation. Once biofilms have grown dense and matured, quorum-sensing can trigger the cells to detach and transition to a planktonic state, leading to dispersal of the cells and potentially tissue colonization when inside the human body. This is governed by the master regulator SmcR, a homolog of LuxR, that senses the cell density with the extracellular concentration of the autoinducer-2 (AI-2) via a signal transduction cascade [62]. SmcR triggers CPS formation (Cluster 2) and the CPS inhibits further biofilm formation and protects the cells in their planktonic state. When inside the mammalian body, CPS plays a critical role in evading the host’s innate immune system by providing antiphagocytic ability and resistance to complement-mediated killing [32]. SmcR expression can also be triggered by mammalian host cells by activating LuxS, an autoinducer-2 synthase [62]. In addition to the CPS and biofilm clusters, genes related to c-di-GMP, quorum-sensing and SmcR were found in several clusters: Clusters 2, 11 and 14 include orthologs earlier shown to be differentially expressed at elevated c-di-GMP levels [40], Cluster 4 and 13 encode diguanylate cyclases that produce c-di-GMP in response to specific environmental signals, Cluster 15 encodes an EAL-domain phosphodiesterase involved in degradation of c-di-GMP, and finally, Cluster 4 includes an ortholog with an SmcR binding site in its promoter region [61], implying its regulation by SmcR.

### Convergent gene losses of clinical-enriched orthologs indicate a shared ecological niche

Most of the clinically enriched orthologs display one of two distinct occurrence patterns among the *V. vulnificus* strains. Of the 128 orthologs, 72 are found in 91-100% of the clinical isolates while being present in 64-90% of the environmental isolates. Meanwhile, 45 are present in 9-22% of the clinical isolates and <1-10% of the environmental isolates (S11A Fig). Ancestral state reconstruction of the orthologs’ presence/absence revealed that all orthologs in the first category had a high likelihood (>96%) of being present at the root of the tree (S11B Fig). This suggests these genes were present in the last common ancestor of *V. vulnificus* but subsequently lost in ancestors of certain clades, primarily those including environmental isolates. In contrast, the 45 orthologs with low prevalence had low likelihood (<20%) of being present in the last common ancestor, indicating they were more likely acquired through multiple independent gene gain events.

Notably, a clade of 19 environmental strains in L2e, along with a clade of four and a clade of one strain in L1, lack a similar set of orthologs that constitute the majority (49–56) of the 72 clinically enriched orthologs that were present in the last common ancestor. This indicates these genes were lost at three independent events in the evolutionary history of *V. vulnificus*, hence an example of convergent evolution. The lost genes are mainly part of co-localisation clusters 3, 14 and 13 on chromosome 1 and clusters 5, 9, 10, 11, 12 on chromosome 2 (Fig 4). Given that these clusters are located at distant loci across both chromosomes, their losses within each clade were unlikely caused by single deletion events, but rather multiple independent deletions, reinforcing the observation of this pattern across different parts of the phylogenetic tree.

Clusters 5, 12, 11, 10 are located (in this order) in a ∼250 kb region on chromosome 2. A possible scenario is that these genes were initially lost in one clade, and subsequently, the whole chromosomal region replaced the corresponding regions in the other two clades through homologous recombination. If this occurred, the remaining genes between the lost co-localisation clusters should exhibit a different phylogeny from the backbone genome tree; the 24 strains with the deletions should form a monophyletic group. To test this hypothesis, we constructed a phylogenetic tree on a concatenation of 82 orthologs located in the regions between clusters 5, 12, 11 and 10. While the tree displayed notable differences from the backbone tree - analogous to variations observed in comparisons of phylogenies for the two chromosomes [22] - the isolates still clustered within the same major lineages as in the backbone tree, including the 24 isolates carrying the deletions (S12 Fig). Therefore, the chromosomal segment replacement scenario appears unlikely, suggesting instead that a more complex evolutionary process resulted in the parallel deletions. The convergent evolution indicates that selection has favored having either all or none of these gene clusters - that include clusters for CPS formation, biofilm formation, pilus synthesis and more - which implies they are functionally connected, and that having or not having them is linked to distinct ecological niches or strategies that *V. vulnificus* can explore in its natural environment. Since these strains lack essential genes for CPS formation, we can be rather sure they are non-pathogenic. It is also noteworthy that these globally distributed strains constitute more than half of the strains that lack essential genes for CPS. Revealing how the niche and environmental drivers of these likely non-pathogenic strains differ from other *V. vulnificus* strains is relevant for proper risk assessment of *V. vulnificus*.

### Potential biomarkers for monitoring of pathogenic *V. vulnificus*

Since *V. vulnificus* infections can rapidly progress into fatal septicemia in susceptible individuals, assays for fast and accurate detection of *V. vulnificus* in environmental (water, sediment) and seafood samples are desirable. Such an assay should specifically detect *V. vulnificus.* Ideally, it should also be specific to pathogenic strains of the species, but more importantly not miss any pathogenic strains, i.e. have a high sensitivity. Genotyping approaches based on genetic variation within individual loci, such as the 16S rRNA, *vcg* (VV1_0734), and *pilF* (VV1_2773) genes have been proposed to distinguish pathogenic from non-pathogenic *V. vulnificus* strains [73–75]. However, the discriminatory power of these loci has been challenged [76,77]. Given that the phylogenetic structure is poorly correlated to clinical vs. environmental isolation source (Fig 2), it seems unlikely (although not impossible) that polymorphism in individual genes would correlate well with pathogenicity. An alternative approach is to base the assay on the presence of a specific gene that occurs in a large fraction of the pathogenic strains of the species while being less frequent in the non-pathogenic ones.

To pinpoint potential biomarkers for identifying pathogenic *V. vulnificus*, we selected a set of “core clinically enriched orthologs” from the pool of the 128 clinically enriched orthologs based on two criteria: the candidates should 1) be present in at least 99.5% of the clinical isolates and 2) exhibit among the highest correlations with clinical isolation source, as determined by the phylogenetic generalized logistic regression (S13 Fig; S4 Table). For eight of these orthologs, we designed primer pairs that were specific to *V. vulnificus* using the degprimer_design and degprimer_specificity pipelines developed as part of this study (see Materials and methods). To assess the efficacy of these primers, we conducted an *in silico* comparative analysis with existing primers for *V. vulnificus* detection (S5 Table). Unlike the existing primers, all of our seven candidate primer pairs demonstrated a match rate exceeding 99.5% in clinical isolates, with a notably greater proportion of matches in clinical compared to environmental strains (83-88%; S6 Table). The fact that it was impossible to design primers matching nearly all clinical strains while at the same time matching nearly none of the environmental ones, is probably reflecting that a substantial fraction of the environmental strains have the genetic potential to be clinical, *i.e.* are pathogenic. We also designed primer pairs that target all *V. vulnificus* strains, with 100% match across all strains (S6 Table). The favorable *in silico* performance of the new primers indicate their potential for surveillance purposes, but follow-up experimental validation is needed.

In conclusion, this study provides new insights into the genomic basis of virulence and ecological differentiation in *Vibrio vulnificus*. By analysing 407 genomes with a phylogeny-aware approach, we identified 128 orthologs significantly enriched in clinical isolates. The organisation of these genes into co-localisation clusters supports the existence of functionally linked pathogenicity modules, including those involved in capsular synthesis, biofilm formation, pilus assembly, and responses to host-like conditions. The observed pattern of convergent gene loss across multiple environmental clades suggests adaptation to non-pathogenic ecological niches, highlighting the evolutionary plasticity of this species. Identifying core clinically enriched orthologs enabled the design of novel, highly specific primers for potential use in monitoring pathogenic *V. vulnificus*. These findings expand our understanding of virulence evolution in this species and lay the groundwork for improved diagnostic and surveillance strategies.

## Materials and methods

### Sampling, isolation, culturing, DNA extraction

Following the salinity gradient of the Baltic Sea, samples from 19 stations were collected from July 25^th^ to September 2^nd^, 2021, covering the German, Estonian, Finnish, Polish, Lithuanian, Swedish, and Danish coasts. From five of these stations (Fig 1), microbial isolates were selected for subsequent genomic analyses. Sampling, isolation and identification of *V. vulnificus* were performed as described in Riedinger et al., (2024) [Citation error]. In short, water and sediment samples were collected from within *Zostera marina* meadows (substation A), as well as from the leaves of *Z. marina* plants. Similarly, water and sediment samples were taken at control stations lacking *Z. marina*, located at distances of 15 m (substation B) and 100 m (substation C) from the meadows’ edges (see interactive S1 File). Sampling was conducted by SCUBA or snorkel divers at water depths ranging from 0.6 to 4.7 m.

*Vibrio* spp. colony forming units (CFUs) and *V. vulnificus* isolates were obtained from water, sediment, and *Z. marina* across six independent replicates. For this, water samples of 50, 100, or 200 μL were (a) directly plated onto Vibrio-selective thiosulfate citrate bile sucrose (TCBS) agar (Merck, Darmstadt, Germany), and (b) aliquots of 2, 5, 10, or 25 mL were filtered onto 0.2 μm PC-filters (Merck-Millipore, Burlington, USA) and placed onto TCBS agar. The sediment samples were homogenized after removing the overlying water; a subsample of approximately 10 g (dry weight determined accurately after lyophilization) was transferred from six sediment samples to sterile 50 mL Falcon tubes, where 40 mL of double 0.2 μm-sterile filtered station water was added. To detach bacteria, five ultrasonic pulses of 10 s at 25% capacity at 5 second intervals using the Bandelin SONOPULS HD 2200.2 (Bandelin, Berlin, Germany) were applied. After subsequent vortexing and settling of the sediment, water samples of 50, 100, or 200 μL were plated on TCBS agar in six biological replicates. For the cultivation of seagrass-associated *Vibrio spp.*, the overlying water was removed; double sterile filtered station water (40 mL) was added to the seagrass samples. To detach bacteria, five ultrasonic pulses of 10s at 25% of capacity at 5-second intervals using the Bandelin SONOPULS HD 2200.2 (Bandelin, Berlin, Germany) were applied to the seagrass samples. After subsequent vortexing and settling of the sediment, between 5 and 20 mL of supernatant was filtered over a PC filter and placed on TCBS-agar in six biological replicates. After 24 h of incubation at 37 °C, CFUs of green colonies were determined for all plates.

To isolate *V. vulnificus*, green colonies were further cultivated on CHROMagar_vibrio™ (Chromagar Ltd. Paris, France) for 24 h at 37 °C, and blue-colored colonies were restreaked on TCBS agar (Carl Roth, Karlsruhe, Germany), CHROMagar (Mast Diagnostica, Reinfeld, Germany), and Columbia sheep blood agar (Oxoid, Basingstoke, UK). DNA of isolates that were green on TCBS, blue on CHROMagar, and confirmed to be pure cultures on blood agar was extracted (DNeasy Blood and Tissue Kit, Qiagen, Hilden, Germany), and purity and concentration were determined using a NanoDrop Spectrophotometer (Thermo Fisher, Waltham, USA). To confirm *V. vulnificus*, the species-specific *vvhA* gene sequence was targeted using multiplex real-time PCR (5’ nuclease assay). In the same assay, an internal amplification control (KOMA) was analysed. Primers and probes for *vvhA* detection were used as described by Messelhäusser et al. (2010) [78].

### DNA sequencing and genome assembly

Shotgun libraries for 84 isolates were prepared using Illumina DNA (Flex) and sequenced together on one Illumina NovaSeq6000 run (2 x 150 bp paired-end reads). Ten of the isolates that obtained too little data were resequenced on one MiSeq (2 x 250 bp paired-end-reads). Reads pre-processing (adapter and low quality bases removal) was carried out with Fastp (params: -- cut_mean_quality 25) [79] and genome assembly with Shovill (v1.1.0, defaults parameters) [80]. Assembly completeness and contamination were evaluated using CheckM [81]. Two genomes were removed because of strain-heterogeneity (>33%), indicating contamination (>83%). The remaining 82 *V. vulnificus* genomes had estimated completeness 100% and contamination 0% and were used for the analyses in this study. We deposited the genomes into GenBank under accession numbers SAMN40604317 - SAMN40604398 under the NCBI BioProject accession no. PRJNA1091677.

### Comparative genomics with PhyloBOTL

The comparative genomic analysis of *V. vulnificus* encompassed 208 clinical isolates (including 43 sourced from the Baltic Sea) and 207 environmental isolates (86 from the Baltic Sea of which 82 were generated in this study), as detailed in S2 Table. Environmental isolates included isolates from water, sediment, and sand sources. Isolates from animal sources were not included. All 407 genomes exhibited estimated completeness >86% (mean 99.92%) and contamination <9.7% (mean 0.17%) (S2 Table). The comparative genomics analysis was executed through the PhyloBOTL pipeline (https://github.com/EnvGen/phyloBOTL). The workflow, illustrated in Fig 5, comprises the following steps (with the parameter settings used in this study indicated):

1. Annotation of genome sequences with Prokka (v1.14.6 with default parameters) [82].
2. Multiple sequence alignment (MSA) of the coding core-genome with ppanggolin (v1.2.105, module ppanggolin msa) [83].
3. Phylogenetic tree inference based on the MSA with maximum likelihood, utilizing IQ-TREE (v2.2.3, parameters: -m GTR+I+G -B 1000 -bnni; and -m 12.12 --seqtype DNA for tree rooting) [84].
4. Identification of orthologous groups of genes with Orthofinder (v2.5.5, parameters: -S diamond -og) [85].
5. Association of orthologs in the genomes to traits of the organisms (in this case, clinical versus environmental source of isolation). We employed the Phylogenetic Generalized Linear Model, utilizing implemented in the R package phylolm [27], which is grounded in an evolutionary model for binary traits, where trait values shift between 0 (environmental isolate) and 1 (clinical isolate) as species evolve along the phylogenetic tree [86]. The R package logistf [87], which implements Firth’s Bias-Reduced Logistic Regression, was employed to achieve precise coefficient estimations when the phylogenetic signal was negligible (alpha > 1628). Firth’s method is recognized as an effective solution to the separation issue in logistic regression [88]. The alpha threshold is defined by the equation log(alpha * T) = 4 [86], where 4 represents the upper limit [27] and T denotes the mean tree tip height. In our analysis, we established the threshold at exp(4) * 0.95 / T, corresponding to 95% of the upper limit values of alpha, which in our case amounts to 1628.
6. Functional annotation of orthologs that are significantly (False Discovery Rate [FDR] adjusted *P*-value < 0.05) enriched or depleted between the traits, using 1) Eggnog mapper (v2.1.12) [89] and the EggNOG database (v5) [90], and 2) mmseqs easy-search (v13.45111, with parameters --max-accept 1 --start-sens 4 -- sens-steps 3 -s 7 -v 1 --min-seq-id 0.9) [91] using Uniref90 [92] proteins with a specific taxonomy affiliation (here, *Vibrionacea*) as reference database.
7. Identification of plasmids and prophages in the genomes using genomad (v1.6.1, with flags --cleanup -- conservative) [93].
8. Grouping the trait-associated orthologs based on proximity in the genomes. Pairs of trait-associated orthologs present within 40 kb distance in at least four genomes were included in the input table to build a network graph, where nodes represent orthologs and edges represent co-occurence in the genomes. The weight of an edge represents the number of genomes in which the ortholog pair co-occurs (within 40 kbp distance). Subsequently, the Leiden graph clustering algorithm, implemented in the R package igraph [94], was applied to obtain the co-localisation clusters. The visualization of the clusters was performed using the R packages gggenomes [95] and ggtree [96], using the script synteny_visual.R of the phyloBOLT pipeline.

**Fig 5.**
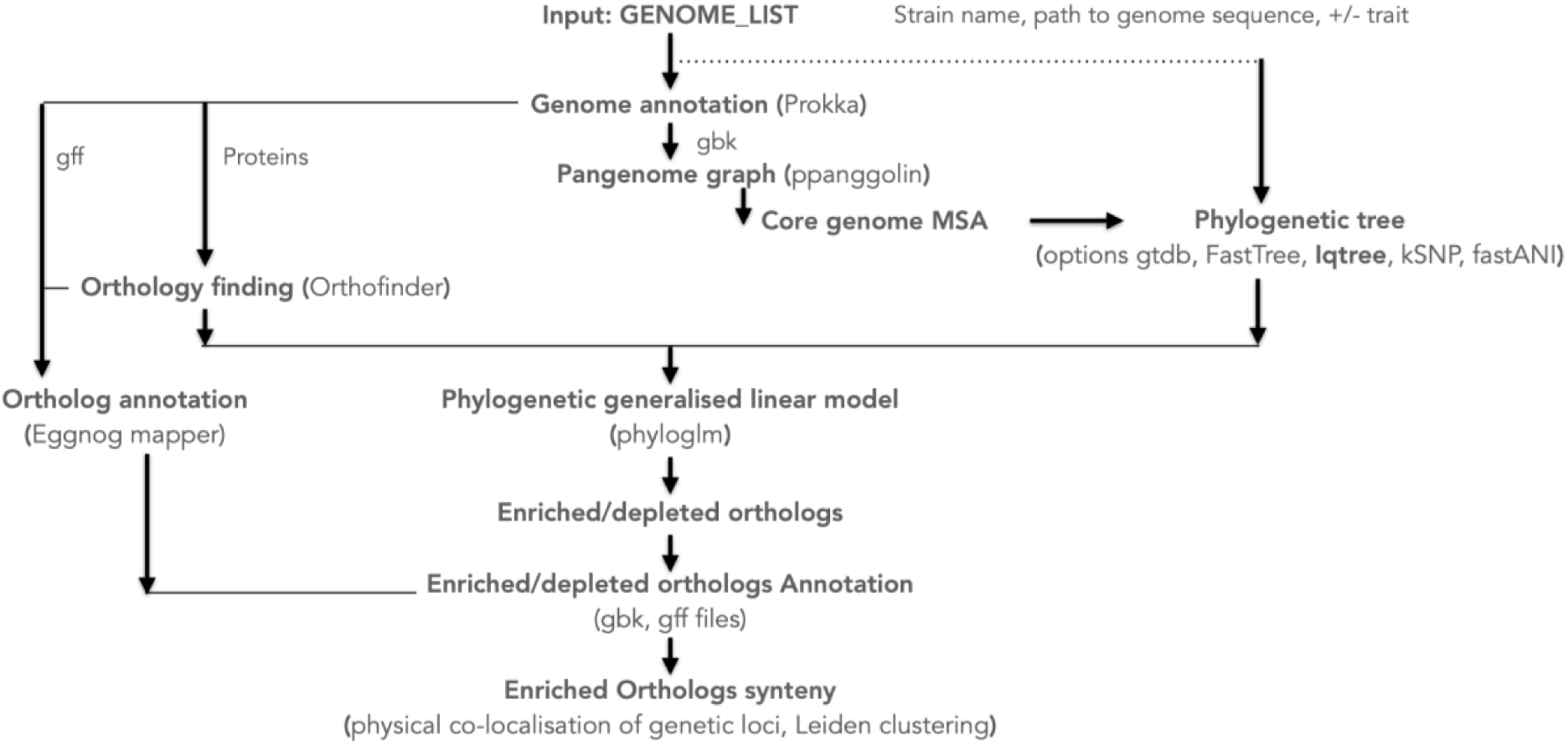
Outline of the PhyloBOTL pipeline workflow.

### Additional bioinformatic analysis

Outside the PhyloBOTL pipeline, we used AMRFinderPlus (version 3.11.26, database version 2023-09-26.1, using the flag --plus) [97] to detect antimicrobial resistance genes in the *V. vulnificus* genomes. We used operon mapper [98] for operon identification in the YJ016, CMCP6 and MO6-24/O complete genomes, and further mapped the genes *de novo* predicted by our pipeline in these three genomes to the corresponding genes in NCBI with Blastx [99], to obtain the original gene names (provided in Table 2). In addition to the automatic gene annotations, we conducted manual annotations of selected clinically enriched orthologs using NCBI BLAST [99] and CDD-search [100], and by literature searches. The tree in Figure 3 was plotted with function plot.phylo in the R package Ape [101]. Ancestral state reconstruction of the orthologs’ presence/absence on the phylogenetic tree was performed with function ace in Ape, using default settings (type = ‘discrete’). For constructing the phylogenetic tree based only on DNA sequences located between the colocalization clusters 5, 10, 11, and 12, the following was performed: On the FORC_009 genome (a complete genome were the gene clusters are present), we identified all orthologs located in the three regions between the clusters (spanning position 294,475 to 531,541 on chromosome 2). Among these orthologs, we selected those that were 1) present in >90% of all strains, 2) never present in >1 copy per genome. The gene sequences of the 82 selected orthologs from each genome (minus potential missing ones) were subsequently concatenated, in the same order.

### Primer design and specificity

The process of identifying candidate biomarkers for *V. vulnificus* pathogenicity involved utilizing enriched orthologous DNA sequences. To design primers, we developed and employed the pipeline Degprimer_design, accessible at https://github.com/envgen/Degprimer_design.

This pipeline integrates several programs, including Snakemake v3.13.3 [102], Muscle v3.8.1551 [103], Degeprime v1.1.0 [104] and Mfeprimer v3.2.3 [105]. Degprimer_design includes the following steps: 1) generation of a multiple sequence alignment of selected orthologous group sequences using Muscle; 2) alignment trimming using TrimAlignment.pl from Degeprime; 3) degenerate primer design using DegePrime.pl from Degeprime for each primer size and degeneracy defined by the user; 4) screening of primers with GC content and coverage above user-defined thresholds; 5) hairpin prediction of screened primers using Mfeprimer; 6) removal of primers forming hairpins; 7) primer-dimer prediction with Mfeprimer and removal of primers forming dimers. 8) Finally, a list of primer pairs is selected based on the user-defined amplicon size, and melting temperature difference between primer pairs below 5°C.

To evaluate primer specificity, we developed and employed the Degprimer_specificity pipeline, accessible at https://github.com/envgen/Degprimer_specificity. This pipeline incorporates Snakemake 3.13.3 [102], Blastn v2.5.0 [99] and Krona v2.7.1 [106]. The Degprimer_specificity pipeline analyzes each primer pair of a list of primer pairs provided by the user. A Blast search is performed against a user-defined database for each non-degenerate version of each degenerate primer. Results are filtered by selecting primer pairs that fulfill the user-defined cutoffs on percent identity (100% in this study) and query coverage, and that generate amplicons in the user-defined size range. *In silico* amplicons are generated and printed to file, together with a Krona file showing the amplicons taxonomic distribution. The output includes a log file (Summary.txt) containing key information, such as the total number of amplicons generated from the target taxon (in this case, *V. vulnificus*) with average amplicon size, as well as the total number of amplicons that are not from the selected taxon. These values can be compared with the values in the Spp_and_strains.txt file, which includes a list and number of species/strains present in the reference database used to check the primer specificity.

Degprimer_specifivity was run on a comprehensive database comprising complete *Vibrio* genomes (51 species, 317 genomes) sourced from The Reference Sequence (RefSeq) project at the National Center for Biotechnology Information (NCBI) [107], alongside 386 draft *V. vulnificus* genomes. This *V. vulnificus* collection includes 43 clinical isolates [10,11] and 82 environmental isolates from the Baltic Sea, as part of the current study. Moreover, we conducted selectivity evaluations on bacteria using a database of complete genomes from Bacteria Refseq [107], encompassing a comprehensive representation of 1097 genera, 2898 species, and 2931 strains.

## Acknowledgements

The authors acknowledge support from the National Genomics Infrastructure in Genomics Production Stockholm funded by Science for Life Laboratory, the Knut and Alice Wallenberg Foundation and the Swedish Research Council, and SNIC/Uppsala Multidisciplinary Center for Advanced Computational Science for assistance with massively parallel sequencing and access to the UPPMAX computational infrastructure.

## Supporting Information

**S1 Fig**. Presence and top 5 most common gene arrangements within co-localization Cluster 3 in the 407 *V. vulnificus* genomes. Beside each gene arrangement, the ID of one genome and the number of genomes containing the arrangement are indicated. Orthologs with blue text are cluster members, grey text non-members. The leaf colors of the tree indicate the presence of different gene arrangements.

**S2 Fig**. Presence and top 5 most common gene arrangements within co-localization Cluster 11 in the 407 *V. vulnificus* genomes. Beside each gene arrangement, the ID of one genome and the number of genomes containing the arrangement are indicated. Orthologs with blue text are cluster members, grey text non-members. The leaf colors of the tree indicate the presence of different gene arrangements.

**S3 Fig**. Presence and top 5 most common gene arrangements within co-localization Cluster 14 in the 407 *V. vulnificus* genomes. Beside each gene arrangement, the ID of one genome and the number of genomes containing the arrangement are indicated. Orthologs with blue text are cluster members, grey text non-members. The leaf colors of the tree indicate the presence of different gene arrangements.

**S4 Fig**. Presence and top 5 most common gene arrangements within co-localization Cluster 4 in the 407 *V. vulnificus* genomes. Beside each gene arrangement, the ID of one genome and the number of genomes containing the arrangement are indicated. Orthologs with blue text are cluster members, grey text non-members. The leaf colors of the tree indicate the presence of different gene arrangements.

**S5 Fig**. Presence and top 5 most common gene arrangements within co-localization Cluster 12 in the 407 *V. vulnificus* genomes. Beside each gene arrangement, the ID of one genome and the number of genomes containing the arrangement are indicated. Orthologs with blue text are cluster members, grey text non-members. The leaf colors of the tree indicate the presence of different gene arrangements.

**S6 Fig**. Presence and top 5 most common gene arrangements within co-localization Cluster 5 in the 407 *V. vulnificus* genomes. Beside each gene arrangement, the ID of one genome and the number of genomes containing the arrangement are indicated. Orthologs with blue text are cluster members, grey text non-members. The leaf colors of the tree indicate the presence of different gene arrangements.

**S7 Fig**. Presence and top 5 most common gene arrangements within co-localization Cluster 10 in the 407 *V. vulnificus* genomes. Beside each gene arrangement, the ID of one genome and the number of genomes containing the arrangement are indicated. Orthologs with blue text are cluster members, grey text non-members. The leaf colors of the tree indicate the presence of different gene arrangements.

**S8 Fig**. Presence and top 5 most common gene arrangements within co-localization Cluster 15 in the 407 *V. vulnificus* genomes. Beside each gene arrangement, the ID of one genome and the number of genomes containing the arrangement are indicated. Orthologs with blue text are cluster members, grey text non-members. The leaf colors of the tree indicate the presence of different gene arrangements.

**S9 Fig**. Presence and top 5 most common gene arrangements within co-localization Cluster 13 in the 407 *V. vulnificus* genomes. Beside each gene arrangement, the ID of one genome and the number of genomes containing the arrangement are indicated. Orthologs with blue text are cluster members, grey text non-members. The leaf colors of the tree indicate the presence of different gene arrangements.

**S10 Fig**. Presence of orthologs related to *V. vulnificus* toxins in the 407 *V. vulnificus* genomes.

**S11 Fig**. Prevalence and ancestral state reconstruction of the 128 clinically enriched orthologs in *V. vulnificus.* A) Prevalence (presence in fraction of genomes) in environmental (x-axis) vs. clinical (y-axis) strains, where each data point is one ortholog. B) Prevalence among all 407 genomes (x-axis) vs. the likelihood that the ortholog was present at the root of the phylogenetic tree (in the last common ancestor of *V. vulnificus*) according to ancestral state reconstruction. The blue and red data points represent orthologs present in 91-100% and 9-22% of the clinical isolates, respectively.

**S12 Fig**. Phylogenetic trees obtained with (A) the full core genome (as in Fig 2 and 4) and (B) with 86 orthologs located between the clinically enriched Clusters 5, 12, 11, 10 in a ∼250 kb region on chromosome 2. The yellow points indicate the strains in which most of the genes of these clusters are missing. These strains are: S3_16, IRLE0056, CECT4608, 1676_80, Vv26, VA_WGS_18029, NV22, BaltVib12-S-24, BaltVib12-S-33, BaltVib12-S-41, VN_0206, BaltVib20-S-8, IRLA0043, VA_WGS_18037, VA_WGS_18038, BaltVib20-S-25, BaltVib20-S-49, BaltVib20-S-5, BaltVib20-W-65, SREL119, IRLE0062, SC9740, IRLE0057 and PV152.

**S13 Fig**. Presence of the core clinically enriched orthologs in the 407 *V. vulnificus* genomes. The orthologs are ordered and colored according to co-localisation clusters.

**S1 File**. Interactive map of sampling locations, including substations, for the *Vibrio vulnificus* isolates.

**S1 Table**. Contextual data for the 82 *V. vulnificus* isolates sequenced in this study.

**S2 Table**. Contextual data and genomic features of the 407 *V. vulnificus* isolates included in this study.

**S3 Table**. Antimicrobial Resistance Genes present in the 407 *V. vulnificus* isolates.

**S4 Table**. Description of the clinically enriched orthologs.

**S5 Table**. Previously published primer pairs for *Vibrio vulnificus* detection.

**S6 Table**. Primer pairs designed in this study for *Vibrio vulnificus* detection.

**S1 Appendix**. Additional information on co-localization Cluster 2, 5, 10, 13 and 15.

**S1 Dataset.** Phylogenetic tree in newick format of the 407 *V. vulnificus* genomes.

**S2 Dataset.** Table with counts of the OGs in the 407 *V. vulnificus* genomes.

**S3 Dataset.** Protein sequences in fasta format for all 13,920 OGs, one representative sequence per OG.

**S4 Dataset.** Gene (DNA) sequences in fasta format for all 13,920 OGs, one representative sequence per OG.

## References

1. Heng S-P, Letchumanan V, Deng C-Y, Ab Mutalib N-S, Khan TM, Chuah L-H, et al. Vibrio vulnificus: An Environmental and Clinical Burden. Front Microbiol. 2017;8: 997.

2. Phillips KE, Satchell KJF. Vibrio vulnificus: From Oyster Colonist to Human Pathogen. PLoS Pathog. 2017;13: e1006053.

3. Hendren N, Sukumar S, Glazer CS. Vibrio vulnificus septic shock due to a contaminated tattoo. BMJ Case Rep. 2017;2017. doi:10.1136/bcr-2017-220199

4. Leng F, Lin S, Wu W, Zhang J, Song J, Zhong M. Epidemiology, pathogenetic mechanism, clinical characteristics, and treatment of Vibrio vulnificus infection: a case report and literature review. Eur J Clin Microbiol Infect Dis. 2019;38: 1999–2004.

5. Elmahdi S, Parveen S, Ossai S, DaSilva LV, Jahncke M, Bowers J, et al. Vibrio parahaemolyticus and Vibrio vulnificus Recovered from Oysters during an Oyster Relay Study. Appl Environ Microbiol. 2018;84. doi:10.1128/AEM.01790-17

6. He R, Zheng W, Long J, Huang Y, Liu C, Wang Q, et al. Vibrio vulnificus meningoencephalitis in a patient with thalassemia and a splenectomy. J Neurovirol. 2019;25: 127–132.

7. Baker-Austin C, Oliver JD. Vibrio vulnificus: new insights into a deadly opportunistic pathogen. Environ Microbiol. 2018;20: 423–430.

8. Jones MK, Oliver JD. Vibrio vulnificus: disease and pathogenesis. Infect Immun. 2009;77: 1723– 1733.

9. Froelich BA, Daines DA. In hot water: effects of climate change on Vibrio-human interactions. Environ Microbiol. 2020;22: 4101–4111.

10. Brehm TT, Berneking L, Sena Martins M, Dupke S, Jacob D, Drechsel O, et al. Heatwave- associated Vibrio infections in Germany, 2018 and 2019. Euro Surveill. 2021;26. doi:10.2807/1560-7917.ES.2021.26.41.2002041

11. Amato E, Riess M, Thomas-Lopez D, Linkevicius M, Pitkänen T, Wołkowicz T, et al. Epidemiological and microbiological investigation of a large increase in vibriosis, northern Europe, 2018. Euro Surveill. 2022;27. doi:10.2807/1560-7917.ES.2022.27.28.2101088

12. Gyraitė G, Kataržytė M, Bučas M, Kalvaitienė G, Kube S, Herlemann DP, et al. Epidemiological and environmental investigation of the “big four” Vibrio species, 1994 to 2021: a Baltic Sea retrospective study. Euro Surveill. 2024;29. doi:10.2807/1560-7917.ES.2024.29.32.2400075

13. Fernández-Juárez V, Riedinger DJ, Gusmao JB, Delgado-Zambrano LF, Coll-García G, Papazachariou V, et al. Temperature, sediment resuspension, and salinity drive the prevalence of Vibrio vulnificus in the coastal Baltic Sea. MBio. 2024;15: e0156924.

14. Riedinger DJ, Fernández-Juárez V, Delgado LF, Sperlea T, Hassenrück C, Herlemann DPR, et al. Control of Vibrio vulnificus proliferation in the Baltic Sea through eutrophication and algal bloom management. Commun Earth Environ. 2024;5. doi:10.1038/s43247-024-01410-x

15. Baker-Austin C, Trinanes JA, Taylor NGH, Hartnell R, Siitonen A, Martinez-Urtaza J. Emerging Vibrio risk at high latitudes in response to ocean warming. Nat Clim Chang. 2013;3: 73–77.

16. Dickerson J Jr, Gooch-Moore J, Jacobs JM, Mott JB. Characteristics of Vibrio vulnificus isolates from clinical and environmental sources. Mol Cell Probes. 2021;56: 101695.

17. Thiaville PC, Bourdage KL, Wright AC, Farrell-Evans M, Garvan CW, Gulig PA. Genotype is correlated with but does not predict virulence of Vibrio vulnificus biotype 1 in subcutaneously inoculated, iron dextran-treated mice. Infect Immun. 2011;79: 1194–1207.

18. Lydon KA, Kinsey T, Le C, Gulig PA, Jones JL. Biochemical and virulence characterization of Vibrio vulnificus isolates from clinical and environmental sources. Front Cell Infect Microbiol. 2021;11: 637019.

19. Sakib SN, Reddi G, Almagro-Moreno S. Environmental role of pathogenic traits in Vibrio cholerae. J Bacteriol. 2018;200: e00795–17.

20. Kim YR, Lee SE, Kim CM, Kim SY, Shin EK, Shin DH, et al. Characterization and pathogenic significance of Vibrio vulnificus antigens preferentially expressed in septicemic patients. Infect Immun. 2003;71: 5461–5471.

21. Bisharat N, Koton Y, Oliver JD. Phylogeography of the marine pathogen, Vibrio vulnificus, revealed the ancestral scenarios of its evolution. Microbiologyopen. 2020;9: e1103.

22. Roig FJ, González-Candelas F, Sanjuán E, Fouz B, Feil EJ, Llorens C, et al. Phylogeny of Vibrio vulnificus from the Analysis of the Core-Genome: Implications for Intra-Species Taxonomy. Front Microbiol. 2017;8: 2613.

23. López-Pérez M, Jayakumar JM, Haro-Moreno JM, Zaragoza-Solas A, Reddi G, Rodriguez-Valera F, et al. Evolutionary model of cluster divergence of the emergent marine pathogen Vibrio vulnificus: From genotype to ecotype. MBio. 2019;10. doi:10.1128/mBio.02852-18

24. Chen C-Y, Wu K-M, Chang Y-C, Chang C-H, Tsai H-C, Liao T-L, et al. Comparative genome analysis of Vibrio vulnificus, a marine pathogen. Genome Res. 2003;13: 2577–2587.

25. López-Pérez M, Jayakumar JM, Grant T-A, Zaragoza-Solas A, Cabello-Yeves PJ, Almagro-Moreno S. Ecological diversification reveals routes of pathogen emergence in endemic Vibrio vulnificus populations. Proc Natl Acad Sci U S A. 2021;118. doi:10.1073/pnas.2103470118

26. Zhang J-X, Yuan Y, Hu Q-H, Jin D-Z, Bai Y, Xin W-W, et al. Identification of potential pathogenic targets and survival strategies of Vibrio vulnificus through population genomics. Front Cell Infect Microbiol. 2023;13: 1254379.

27. Ho L si T, Ané C. A linear-time algorithm for Gaussian and non-Gaussian trait evolution models. Syst Biol. 2014;63: 397–408.

28. Liu B, Zheng D, Zhou S, Chen L, Yang J. VFDB 2022: a general classification scheme for bacterial virulence factors. Nucleic Acids Res. 2022;50: D912–D917.

29. Overbeek R, Fonstein M, D’Souza M, Pusch GD, Maltsev N. The use of gene clusters to infer functional coupling. Proc Natl Acad Sci U S A. 1999;96: 2896–2901.

30. Dandekar T, Snel B, Huynen M, Bork P. Conservation of gene order: a fingerprint of proteins that physically interact. Trends Biochem Sci. 1998;23: 324–328.

31. Wang X, Ma Q. Wzb of Vibrio vulnificus represents a new group of low-molecular-weight protein tyrosine phosphatases with a unique insertion in the W-loop. J Biol Chem. 2021;296. doi:10.1016/j.jbc.2021.100280

32. Pettis GS, Mukerji AS. Structure, Function, and Regulation of the Essential Virulence Factor Capsular Polysaccharide of Vibrio vulnificus. Int J Mol Sci. 2020;21: 3259.

33. Linkous DA, Oliver JD. Pathogenesis of Vibrio vulnificus. FEMS Microbiol Lett. 1999;174: 207– 214.

34. Hayat U, Reddy GP, Bush CA, Johnson JA, Wright AC, Morris JG. Capsular Types of Vibrio vulnificus: An Analysis of Strains from Clinical and Environmental Sources. J Infect Dis. 1993;168: 758–762.

35. Hilton Tamara, Rosche Tom, Froelich Brett, Smith Benjamin, Oliver James. Capsular Polysaccharide Phase Variation in Vibrio vulnificus. Appl Environ Microbiol. 2006;72: 6986–6993.

36. Wright A C, Simpson L M, Oliver J D, Morris J G. Phenotypic evaluation of acapsular transposon mutants of Vibrio vulnificus. Infect Immun. 1990;58: 1769–1773.

37. Nakhamchik Alina, Wilde Caroline, Rowe-Magnus Dean A. Cyclic-di-GMP Regulates Extracellular Polysaccharide Production, Biofilm Formation, and Rugose Colony Development by Vibrio vulnificus. Appl Environ Microbiol. 2008;74: 4199–4209.

38. Joseph LA, Wright AC. Expression of Vibrio vulnificus capsular polysaccharide inhibits biofilm formation. J Bacteriol. 2004;186: 889–893.

39. Wright Anita C., Powell Jan L., Kaper James B., Morris J. Glenn. Identification of a Group 1-Like Capsular Polysaccharide Operon for Vibrio vulnificus. Infect Immun. 2001;69: 6893–6901.

40. Chodur DM, Rowe-Magnus DA. Complex Control of a Genomic Island Governing Biofilm and Rugose Colony Development in Vibrio vulnificus. J Bacteriol. 2018;200. doi:10.1128/JB.00190-18

41. Park JH, Jo Y, Jang SY, Kwon H, Irie Y, Parsek MR, et al. The cabABC Operon Essential for Biofilm and Rugose Colony Development in Vibrio vulnificus. PLoS Pathog. 2015;11: e1005192.

42. Park JH, Lee B, Jo Y, Choi SH. Role of extracellular matrix protein CabA in resistance of Vibrio vulnificus biofilms to decontamination strategies. Int J Food Microbiol. 2016;236: 123–129.

43. Hwang S-H, Park JH, Lee B, Choi SH. A Regulatory Network Controls cabABC Expression Leading to Biofilm and Rugose Colony Development in Vibrio vulnificus. Front Microbiol. 2019;10: 3063.

44. Hwang S-H, Im H, Choi SH. A Master Regulator BrpR Coordinates the Expression of Multiple Loci for Robust Biofilm and Rugose Colony Development in Vibrio vulnificus. Front Microbiol. 2021;12: 679854.

45. Chodur DM, Coulter P, Isaacs J, Pu M, Fernandez N, Waters CM, et al. Environmental Calcium Initiates a Feed-Forward Signaling Circuit That Regulates Biofilm Formation and Rugosity in Vibrio vulnificus. MBio. 2018;9. doi:10.1128/mBio.01377-18

46. Choi G, Choi SH. Complex regulatory networks of virulence factors in Vibrio vulnificus. Trends Microbiol. 2022;30: 1205–1216.

47. Lee H, Im H, Hwang S-H, Ko D, Choi SH. Two novel genes identified by large-scale transcriptomic analysis are essential for biofilm and rugose colony development of Vibrio vulnificus. PLoS Pathog. 2023;19: e1011064.

48. Nuccio S-P, Bäumler AJ. Evolution of the chaperone/usher assembly pathway: fimbrial classification goes Greek. Microbiol Mol Biol Rev. 2007;71: 551–575.

49. Pakharukova N, Malmi H, Tuittila M, Dahlberg T, Ghosal D, Chang Y-W, et al. Archaic chaperone- usher pili self-secrete into superelastic zigzag springs. Nature. 2022;609: 335–340.

50. Tomaras AP, Dorsey CW, Edelmann RE, Actis LA. Attachment to and biofilm formation on abiotic surfaces by Acinetobacter baumannii: involvement of a novel chaperone-usher pili assembly system. Microbiology. 2003;149: 3473–3484.

51. Giraud C, Bernard CS, Calderon V, Yang L, Filloux A, Molin S, et al. The PprA-PprB two-component system activates CupE, the first non-archetypal Pseudomonas aeruginosa chaperone- usher pathway system assembling fimbriae. Environ Microbiol. 2011;13: 666–683.

52. Paranjpye RN, Strom MS. A Vibrio vulnificus type IV pilin contributes to biofilm formation, adherence to epithelial cells, and virulence. Infect Immun. 2005;73: 1411–1422.

53. Pu M, Rowe-Magnus DA. A Tad pilus promotes the establishment and resistance of Vibrio vulnificus biofilms to mechanical clearance. NPJ Biofilms Microbiomes. 2018;4: 10.

54. Salomon D, Kinch LN, Trudgian DC, Guo X, Klimko JA, Grishin NV, et al. Marker for type VI secretion system effectors. Proc Natl Acad Sci U S A. 2014;111: 9271–9276.

55. Shneider MM, Buth SA, Ho BT, Basler M, Mekalanos JJ, Leiman PG. PAAR-repeat proteins sharpen and diversify the type VI secretion system spike. Nature. 2013;500: 350–353.

56. Russell AB, Peterson SB, Mougous J. Type VI secretion system effectors: poisons with a purpose. Nat Rev Microbiol. 2014;12: 137–148.

57. Wiener M, Freymann D, Ghosh P, Stroud RM. Crystal structure of colicin Ia. Nature. 1997;385: 461–464.

58. Dar Y, Salomon D, Bosis E. The Antibacterial and Anti-Eukaryotic Type VI Secretion System MIX-Effector Repertoire in Vibrionaceae. Mar Drugs. 2018;16. doi:10.3390/md16110433

59. Church SR, Lux T, Baker-Austin C, Buddington SP, Michell SL. Vibrio vulnificus Type 6 Secretion System 1 Contains Anti-Bacterial Properties. PLoS One. 2016;11: e0165500.

60. Hubert CL, Michell SL. A universal oyster infection model demonstrates that Vibrio vulnificus Type 6 secretion systems have antibacterial activity in vivo. Environ Microbiol. 2020;22: 4381– 4393.

61. Lee DH, Jeong HS, Jeong HG, Kim KM, Kim H, Choi SH. A consensus sequence for binding of SmcR, a Vibrio vulnificus LuxR homologue, and genome-wide identification of the SmcR regulon. J Biol Chem. 2008;283: 23610–23618.

62. Kim SM, Park JH, Lee HS, Kim WB, Ryu JM, Han HJ, et al. LuxR homologue SmcR is essential for Vibrio vulnificus pathogenesis and biofilm detachment, and its expression is induced by host cells. Infect Immun. 2013;81: 3721–3730.

63. Chou L-F, Peng H-L, Yang Y-C, Kuo M-C, Chang H-Y. Localization and characterization of VVA0331, a 489-kDa RTX-like protein, in Vibrio vulnificus YJ016. Arch Microbiol. 2009;191: 441–450.

64. Gulig PA, Bourdage KL, Starks AM. Molecular pathogenesis ofVibrio vulniWcus. J Microbiol. 2005;43: 118–131.

65. Nair AV, Singh A, Rajmani RS, Chakravortty D. Salmonella Typhimurium employs spermidine to exert protection against ROS-mediated cytotoxicity and rewires host polyamine metabolism to ameliorate its survival in macrophages. Redox Biol. 2024;72: 103151.

66. Williams TC, Blackman ER, Morrison SS, Gibas CJ, Oliver JD. Transcriptome sequencing reveals the virulence and environmental genetic programs of Vibrio vulnificus exposed to host and estuarine conditions. PLoS One. 2014;9: e114376.

67. McAdow M, Kim HK, Dedent AC, Hendrickx APA, Schneewind O, Missiakas DM. Preventing Staphylococcus aureus sepsis through the inhibition of its agglutination in blood. PLoS Pathog. 2011;7: e1002307.

68. Kim BS. The Modes of Action of MARTX Toxin Effector Domains. Toxins . 2018;10. doi:10.3390/toxins10120507

69. Jang KK, Lee Z-W, Kim B, Jung YH, Han HJ, Kim MH, et al. Identification and characterization of Vibrio vulnificus plpA encoding a phospholipase A2 essential for pathogenesis. J Biol Chem. 2017;292: 17129–17143.

70. Yuan Y, Feng Z, Wang J. Vibrio vulnificus Hemolysin: Biological Activity, Regulation of vvhA Expression, and Role in Pathogenesis. Front Immunol. 2020;11: 599439.

71. Miyoshi S-I. Vibrio vulnificus infection and metalloprotease. J Dermatol. 2006;33: 589–595.

72. Ashrafudoulla M, Mizan MFR, Park SH, Ha S-D. Current and future perspectives for controlling Vibrio biofilms in the seafood industry: a comprehensive review. Crit Rev Food Sci Nutr. 2021;61: 1827–1851.

73. Nilsson William B., Paranjype Rohinee N., DePaola Angelo, Strom Mark S. Sequence Polymorphism of the 16S rRNA Gene of Vibrio vulnificus Is a Possible Indicator of Strain Virulence. J Clin Microbiol. 2003;41: 442–446.

74. Rosche TM, Yano Y, Oliver JD. A rapid and simple PCR analysis indicates there are two subgroups of Vibrio vulnificus which correlate with clinical or environmental isolation. Microbiol Immunol. 2005;49: 381–389.

75. Roig FJ, Sanjuán E, Llorens A, Amaro C. pilF Polymorphism-based PCR to distinguish vibrio vulnificus strains potentially dangerous to public health. Appl Environ Microbiol. 2010;76: 1328– 1333.

76. Sanjuán E, Fouz B, Oliver JD, Amaro C. Evaluation of genotypic and phenotypic methods to distinguish clinical from environmental Vibrio vulnificus strains. Appl Environ Microbiol. 2009;75: 1604–1613.

77. Bier N, Bechlars S, Diescher S, Klein F, Hauk G, Duty O, et al. Genotypic diversity and virulence characteristics of clinical and environmental Vibrio vulnificus isolates from the Baltic Sea region. Appl Environ Microbiol. 2013;79: 3570–3581.

78. Messelhäusser U, Colditz J, Thärigen D, Kleih W, Höller C, Busch U. Detection and differentiation of Vibrio spp. in seafood and fish samples with cultural and molecular methods. Int J Food Microbiol. 2010;142: 360–364.

79. Chen S, Zhou Y, Chen Y, Gu J. fastp: an ultra-fast all-in-one FASTQ preprocessor. Bioinformatics. 2018;34: i884–i890.

80. Seemann T. shovill: ⚡♠ Assemble bacterial isolate genomes from Illumina paired-end reads. Github; Available: https://github.com/tseemann/shovill

81. Parks DH, Imelfort M, Skennerton CT, Hugenholtz P, Tyson GW. CheckM: assessing the quality of microbial genomes recovered from isolates, single cells, and metagenomes. Genome Res. 2015;25: 1043–1055.

82. Seemann T. Prokka: rapid prokaryotic genome annotation. Bioinformatics. 2014;30: 2068–2069.

83. Gautreau G, Bazin A, Gachet M, Planel R, Burlot L, Dubois M, et al. Correction: PPanGGOLiN: Depicting microbial diversity via a partitioned pangenome graph. PLoS Comput Biol. 2021;17: e1009687.

84. Minh BQ, Schmidt HA, Chernomor O, Schrempf D, Woodhams MD, von Haeseler A, et al. Corrigendum to: IQ-TREE 2: New Models and Efficient Methods for Phylogenetic Inference in the Genomic Era. Mol Biol Evol. 2020;37: 2461.

85. Emms DM, Kelly S. OrthoFinder: phylogenetic orthology inference for comparative genomics. Genome Biol. 2019;20: 238.

86. Ives AR, Garland T Jr. Phylogenetic logistic regression for binary dependent variables. Syst Biol. 2010;59: 9–26.

87. Heinze G, Ploner M, Jiricka L, Steiner G. logistf: Firth’s Bias-Reduced Logistic Regression. R package version 1.26.0. Available: https://CRAN.R-project.org/package=logistf

88. Heinze G, Schemper M. A solution to the problem of separation in logistic regression. Stat Med. 2002;21: 2409–2419.

89. Cantalapiedra CP, Hernández-Plaza A, Letunic I, Bork P, Huerta-Cepas J. eggNOG-mapper v2: Functional Annotation, Orthology Assignments, and Domain Prediction at the Metagenomic Scale. Mol Biol Evol. 2021;38: 5825–5829.

90. Huerta-Cepas J, Szklarczyk D, Heller D, Hernández-Plaza A, Forslund SK, Cook H, et al. eggNOG 5.0: a hierarchical, functionally and phylogenetically annotated orthology resource based on 5090 organisms and 2502 viruses. Nucleic Acids Res. 2019;47: D309–D314.

91. Hauser M, Steinegger M, Söding J. MMseqs software suite for fast and deep clustering and searching of large protein sequence sets. Bioinformatics. 2016;32: 1323–1330.

92. Suzek BE, Huang H, McGarvey P, Mazumder R, Wu CH. UniRef: comprehensive and non-redundant UniProt reference clusters. Bioinformatics. 2007;23: 1282–1288.

93. Fast and accurate identification of plasmids and viruses in sequencing data using geNomad. Nat Biotechnol. 2023. doi:10.1038/s41587-023-01982-7

94. Csárdi G, Horvát S, Müller K, Nepusz T, Noom D, Salmon M, et al. igraph: network analysis and visualization. R package version 1.2. 4.1. 2019.

95. Hackl T, Ankenbrand MJ. gggenomes: a grammar of graphics for comparative genomics. R package version 09.

96. Yu G. Data Integration, Manipulation and Visualization of Phylogenetic Trees. CRC Press; 2022.

97. Feldgarden M, Brover V, Gonzalez-Escalona N, Frye JG, Haendiges J, Haft DH, et al. AMRFinderPlus and the Reference Gene Catalog facilitate examination of the genomic links among antimicrobial resistance, stress response, and virulence. Sci Rep. 2021;11: 12728.

98. Taboada B, Estrada K, Ciria R, Merino E. Operon-mapper: a web server for precise operon identification in bacterial and archaeal genomes. Bioinformatics. 2018;34: 4118–4120.

99. Camacho C, Coulouris G, Avagyan V, Ma N, Papadopoulos J, Bealer K, et al. BLAST+: architecture and applications. BMC Bioinformatics. 2009;10: 421.

100. Wang J, Chitsaz F, Derbyshire MK, Gonzales NR, Gwadz M, Lu S, et al. The conserved domain database in 2023. Nucleic Acids Res. 2023;51: D384–D388.

101. Paradis E, Schliep K. ape 5.0: an environment for modern phylogenetics and evolutionary analyses in R. Bioinformatics. 2019;35: 526–528.

102. Köster J, Rahmann S. Snakemake—a scalable bioinformatics workflow engine. Bioinformatics. 2012. Available: https://academic.oup.com/bioinformatics/article-abstract/28/19/2520/290322

103. Edgar RC. MUSCLE: multiple sequence alignment with high accuracy and high throughput. Nucleic Acids Res. 2004;32: 1792–1797.

104. Hugerth LW, Wefer HA, Lundin S, Jakobsson HE, Lindberg M, Rodin S, et al. DegePrime, a program for degenerate primer design for broad-taxonomic-range PCR in microbial ecology studies. Appl Environ Microbiol. 2014;80: 5116–5123.

105. Wang K, Li H, Xu Y, Shao Q, Yi J, Wang R, et al. MFEprimer-3.0: quality control for PCR primers. Nucleic Acids Res. 2019;47: W610–W613.

106. Ondov BD, Bergman NH, Phillippy AM. Interactive metagenomic visualization in a Web browser. BMC Bioinformatics. 2011;12: 385.

107. Haft DH, DiCuccio M, Badretdin A, Brover V, Chetvernin V, O’Neill K, et al. RefSeq: an update on prokaryotic genome annotation and curation. Nucleic Acids Res. 2018;46: D851–D860.

